# Strain-dependent effects on neurobehavioral and seizure phenotypes in *Scn2a^K1422E^* mice

**DOI:** 10.1101/2023.06.06.543929

**Authors:** Dennis M. Echevarria-Cooper, Nicole A. Hawkins, Jennifer A. Kearney

## Abstract

Pathogenic variants in *SCN2A* are associated with a range of neurodevelopmental disorders (NDD). Despite being largely monogenic, *SCN2A*-related NDD show considerable phenotypic variation and complex genotype-phenotype correlations. Genetic modifiers can contribute to variability in disease phenotypes associated with rare driver mutations. Accordingly, different genetic backgrounds across inbred rodent strains have been shown to influence disease-related phenotypes, including those associated with *SCN2A*-related NDD. Recently, we developed a mouse model of the variant *SCN2A*-p.K1422E that was maintained as an isogenic line on the C57BL/6J (B6) strain. Our initial characterization of NDD phenotypes in heterozygous *Scn2a^K1422E^* mice revealed alterations in anxiety-related behavior and seizure susceptibility. To determine if background strain affects phenotype severity in the *Scn2a^K1422E^* mouse model, phenotypes of mice on B6 and [DBA/2JxB6]F1 hybrid (F1D2) strains were compared.

Convergent evidence from neurobehavioral assays demonstrated lower anxiety-like behavior in *Scn2a^K1422E^* mice compared to wild-type and further suggested that this effect is more pronounced on the B6 background compared to the F1D2 background. Although there were no strain-dependent differences in occurrence of rare spontaneous seizures, response to the chemoconvulsant kainic acid revealed differences in seizure generalization and lethality risk, with variation based on strain and sex. Continued examination of strain-dependent effects in the *Scn2a^K1422E^* mouse model could reveal genetic backgrounds with unique susceptibility profiles that would be relevant for future studies on specific traits and enable the identification of highly penetrant phenotypes and modifier genes that could provide clues about the primary pathogenic mechanism of the K1422E variant.

## Introduction

*SCN2A*, encoding the Na_V_1.2 voltage-gated sodium channel, has been established as a major genetic driver of neurodevelopmental disorders (NDD)^1^. Efforts to better understand the complex spectrum of *SCN2A*-related NDD have been aimed at cataloging pathogenic variants based on their association with distinct biophysical defects in channel function. This process has established a general framework wherein missense variants with gain-of-function effects have been associated with severe, early-onset epilepsy phenotypes while missense and nonsense variants with loss-of function effects have been associated with autism spectrum disorder (ASD) and intellectual disability^2–6^. Although the biophysical properties of mutant channels likely contribute to phenotype expressivity, recurrent and inherited variants in *SCN2A* show wide phenotypic heterogeneity, even among individuals with the same variant^2, 3, 7^. This suggests that other factors interact with *SCN2A* variants to influence neuropathological phenotypes. It is becoming increasingly evident that background genetic variation (i.e., modifiers) can affect the phenotypic properties of presumed monogenic disorders, but this can be difficult to study in the context of rare diseases with small patient populations^8–14^.

Inbred rodent populations (strains) can have mixed effects when it comes to modeling human disease. On the one hand, the isogenic nature of these animals makes them attractive for use as biological replicates^15^. On the other hand, the genetic background of particular strains may interact with the variant gene of interest in a way that biases the analysis of disease-related phenotypes^16^. However, these properties can be leveraged to study the effect of genetic modifiers in rodent models of disease. Accordingly, different genetic backgrounds across rodent strains have been shown to influence disease-related phenotypes, including seizure susceptibility^16–26^. For example, it has been demonstrated that genetic background dramatically influences the epilepsy phenotype described in the *Scn2a^Q^*^54^ mouse model. *Scn2a^Q^*^54^ mice on a C57BL/6J (B6) background displayed adult-onset seizures and a mild survival deficit (>75% survival at 6 months). In contrast, *Scn2a^Q^*^54^ mice on a [B6 x SJL/J]F1 background displayed juvenile-onset seizures and a more severe survival deficit (25% survival at 6 months)^17, 18^. Studies on the effect of genetic background on seizure susceptibility can serve as the basis for identification of epilepsy modifier loci and/or genes. Using both classical genetic mapping and candidate gene approaches, a number of genes, including *Hlf, Kcnv2, Cacna1g, Kcnq2,* and *Scn1a* were identified as modifiers of *Scn2a^Q^*^54, 18, 24, 27–34^. Identifying modifier genes can enhance our understanding of disease mechanisms and support the development of targeted therapeutic strategies.

Previously, we established a mouse model of the *SCN2A* variant p.K1422E on the B6 background and showed that heterozygous *Scn2a^K1422E^* (*Scn2a^E/+^*) mice exhibit NDD-related phenotypes, including alterations in anxiety-like behavior and seizure susceptibility^35^. Given the demonstrated effects of genetic background in other epilepsy models like *Scn2a^Q^*^54^, we hypothesized that background strain would affect phenotype severity in the *Scn2a^K1422E^*mouse model. To address this hypothesis, we crossed B6.*Scn2a^E/+^*males to inbred DBA/2J (D2) females to generate F1 [D2xB6] hybrid offspring (F1D2). It was previously shown that D2 mice exhibit a high rate of spontaneous spike-wave discharges associated with behavioral arrest, and are more susceptible to focal and generalized seizures induced by kainic acid, as well as to generalized seizures induced by maximal electroshock^36–40^. We evaluated anxiety-like behavior and seizure susceptibility in wild-type (WT) and *Scn2a^E/+^* mice on the F1D2 background compared to mice on the B6 background. Convergent evidence from neurobehavioral assays demonstrates lower anxiety-like behavior in *Scn2a^E/+^* mice compared to WT and further suggests the effect is more pronounced on the B6 background compared to the F1D2 background.

F1D2*.Scn2a^E/+^* mice also exhibited rare spontaneous seizures similar to those previously observed in B6*.Scn2a^E/+^* mice^35^. Following administration of the chemoconvulsant kainic acid (KA), *Scn2a^E/+^* mice were less susceptible to seizure generalization but had increased risk of lethality relative to WT, with variation based on strain and sex. These data suggest that background strain does affect phenotype severity in the *Scn2a^K1422E^* mouse model.

## Materials and Methods

### Mice

The line *Scn2a^em1Kea^* (RRID:MMRRC_069700-UCD), is maintained as an isogenic strain on C57BL/6J (#000664, Jackson Laboratory, Bar Harbor, ME). B6.*Scn2a^E/+^* males were crossed to inbred DBA/2J females (#000671, Jackson Laboratory) to generate F1 [D2xB6] hybrid offspring. All animal care and experimental procedures were approved by the Northwestern University Animal Care and Use Committees in accordance with the National Institutes of Health Guide for the Care and Use of Laboratory Animals. Principles outlined in the ARRIVE (Animal Research: Reporting of *in vivo* Experiments) guideline were considered when planning experiments^41^.

### Neurobehavioral assays

B6 and F1D2 WT and *Scn2a^E/+^* mice of both sexes were tested at 8-12 weeks of age. Male and female mice were tested on separate days. For all experiments, mice were acclimated in the behavior suite with white noise for 1 h prior to behavioral testing. At the end of each procedure, mice were placed into their home cage with their original littermates.

Behavioral testing was performed by experimenters blinded to genotype. The same cohort of mice was evaluated first in the zero maze and then in the open field, with a two-day interval between assays. For all measures male and females were considered separately. Statistical comparisons between groups were made using ordinary two-way ANOVA with Sidak’s post-hoc comparisons (**Table 1**).

**Table 1.** Statistical comparisons.

| Figure | Comparison | Test | Value | Post Hoc |
| --- | --- | --- | --- | --- |
| **1** | Zero Maze – Open arm time<br>Male (A) | Two-way ANOVA | F(1,37)=47.46, p<0.0001<br>(Main Effect: Strain)<br>F(1,37)=9.877, p=0.0033<br>(Main Effect: Genotype) | Sidak's |
|  | Zero Maze – Open arm time<br>Female (A) | Two-way ANOVA | F(1,40)=23.26, p<0.0001<br>(Strain × Genotype)<br>F(1,40)=242.4, p<0.0001<br>(Main Effect: Strain)<br>F(1,40)=74.02, p<0.0001<br>(Main Effect: Genotype) | Sidak's |
|  | Zero Maze – Distance traveled<br>Male (B) | Two-way ANOVA | F(1,37)=40.02, p<0.0001<br>(Main Effect: Strain)<br>F(1,37)=8.686, p=0.0055<br>(Main Effect: Genotype) | Sidak's |
|  | Zero Maze – Distance traveled<br>Female (B) | Two-way ANOVA | F(1,40)=107.8, p<0.0001<br>(Main Effect: Strain)<br>F(1,40)=11.89, p=0.0013<br>(Main Effect: Genotype) | Sidak's |
|  | Open Field – Center time<br>Male (C) | Two-way ANOVA | F(1,45)=13.52, p=0.0006<br>(Strain × Genotype)<br>F(1,45)=7.189, p=0.0102<br>(Main Effect: Strain) | Sidak's |
|  | Open Field – Center time<br>Female (C) | Two-way ANOVA | F(1,43)=19.61, p<0.0001<br>(Main Effect: Strain) | Sidak's |
|  | Open Field – Distance traveled<br>Male (D) | Two-way ANOVA | F(1,45)=32.78, p<0.0001<br>(Main Effect: Strain) | Sidak's |
|  | Open Field – Distance traveled<br>Female (D) | Two-way ANOVA | F(1,43)=57.73, p<0.0001<br>(Main Effect: Strain) | Sidak's |
| **4** | KA stage 3 – B6 E/+<br>SEX (A) | Log-rank Mantel-Cox test | p=0.0073 | n/a |
|  | KA stage 3 – F1D2 Male<br>GENOTYPE (A) | Log-rank Mantel-Cox test | p=0.0240 | n/a |
|  | KA stage 6 – B6 E/+<br>SEX (B) | Log-rank Mantel-Cox test | p=0.0107 | n/a |
|  | KA stage 6 – B6 Male<br>GENOTYPE (B) | Log-rank Mantel-Cox test | p=0.0003 | n/a |
|  | KA stage 6 – F1D2 Male<br>GENOTYPE (B) | Log-rank Mantel-Cox test | p<0.0001 | n/a |
|  | KA stage 6 – F1D2 Female<br>GENOTYPE (B) | Log-rank Mantel-Cox test | p<0.0001 | n/a |
|  | KA stage 6 – E/+ Female<br>STRAIN (B) | Log-rank Mantel-Cox test | p=0.0008 | n/a |
|  | KA stage 7 (lethality) – B6 E/+<br>SEX (C) | Log-rank Mantel-Cox test | p=0.0077 | n/a |
|  | KA stage 7 (lethality) – B6 Male<br>GENOTYPE (C) | Log-rank Mantel-Cox test | p=0.0467 | n/a |
|  | KA stage 7 (lethality) – WT Male<br>STRAIN (C) | Log-rank Mantel-Cox test | p=0.0001 | n/a |
|  | KA stage 7 (lethality) – WT Female<br>STRAIN (C) | Log-rank Mantel-Cox test | p=0.0002 | n/a |
|  | KA stage 7 (lethality) – E/+ Female<br>STRAIN (C) | Log-rank Mantel-Cox test | p=0.0006 | n/a |

### Zero maze

Mice were evaluated for anxiety-related behavior in an elevated zero maze. The maze consists of an annular platform (diameter 46 cm; elevation 50 cm) divided into equally sized quadrants, alternating between open and enclosed (wall height 17 cm). This configuration lacks the ambiguous center region associated with the elevated plus maze^42^. The open and closed arms were illuminated to similar levels (30 lx). Individual mice were placed near an enclosed arm of the maze and allowed to freely explore for 5 min. Limelight software (Actimetrics, Wilmette, IL, USA) was used to video record each trial. Ethovison XT software (Noldus, Leesberg, VA, USA) was used to track the position of the mouse, and calculate distance traveled, mean velocity and time spent in closed or open arms (*n* = 6–14 per strain, genotype, and sex).

Trials where mice exited the maze were excluded from analysis. Exclusions by group were as follows: B6.*Scn2a^E/+^* males – 7 of 13, B6.*Scn2a^E/+^* females – 5 of 15, F1D2.*Scn2a^E/+^*males – 2 of 11; all other groups had no exclusions. The proportion of B6.*Scn2a^E/+^*males excluded due to maze exit was significantly greater than B6.WT males (p*=*0.0052, Fisher’s exact test), while the proportion of B6.*Scn2a^E/+^* females excluded was significantly greater than F1D2.*Scn2a^E/+^* females (p*=*0.0421, Fisher’s exact test).

### Open field

Mice were evaluated for baseline activity and anxiety-related behavior in an open field. Individual mice were placed in the center of an open field arena (46 × 46 cm, illuminance 70 lx) and allowed to freely explore for 30 min. Limelight software was used to video record each trial. Ethovison XT software was used to track the position of the mouse, and calculate total distance traveled, mean velocity and time spent in the periphery (9 cm from wall) and center (28 × 28 cm) (*n* = 9–15 per strain, genotype, and sex).

### Video-EEG monitoring

Male and female F1D2.WT and F1D2.*Scn2a^E/+^* mice were implanted with prefabricated 3-channel EEG headmounts (Pinnacle Technology, Lawrence, KS, USA) at P28-30 days of age. Briefly, mice were anesthetized with ketamine/xylazine and place in a stereotaxic frame. Headmounts with four stainless steel screws that serve as cortical surface electrodes were affixed to the skull with glass ionomer cement. Anterior screw electrodes were 0.5–1 mm anterior to bregma and 1 mm lateral from the midline. Posterior screws were 4.5–5 mm posterior to bregma and 1 mm lateral from the midline. EEG1 represents recordings from right posterior to left posterior (interelectrode distance ≈ 2 mm). EEG2 represents recordings from right anterior to left posterior (interelectrode distance ≈ 5 mm). The left anterior screw served as the ground connection. Following at least 5 days of recovery, tethered EEG and video data were continuously collected from freely moving mice with Sirenia acquisition software (Pinnacle Technology) using a sampling rate of 400 Hz as previously described^43^. Between 168-240 h of EEG data were acquired from each subject (F1D2.WT: *n* = 9 mice, 4-5 weeks of age; F1D2.*Scn2a^E/+^*: *n* = 14 mice, 4–7 weeks of age). Raw data was notch filtered with a 1 Hz window around 60 and 120 Hz prior to analysis. Video-EEG records were manually reviewed with Sirenia software by three independent reviewers blinded to genotype. Spontaneous seizures were defined as isolated events with an amplitude of ≥3 times baseline, duration of ≥10 s and that show evolution in power and amplitude. Data from five subjects with high baseline artifact or low signal were excluded from analysis.

### Kainic acid seizure induction

Susceptibility to seizures induced by the chemoconvulsant kainic acid (abbreviated as KA, Cat #0222, Tocris Bioscience, Minneapolis, MN) was assessed in B6 and F1D2 WT and *Scn2a^E/+^*mice of both sexes P41-44 days of age. KA dissolved in saline to a working concentration of 2.5 mg/mL was administered by IP injection (25 mg/kg). Mice were placed in clean cages and video recorded (side view) for 2 hours. Videos were scored offline by two reviewers blinded to genotype using a modified Racine scale^44^ (1– behavioral arrest; 2– forelimb and/or Straub tail, facial automatisms; 3– automatisms, including repetitive scratching, circling, forelimb clonus without falling; 4– forelimb clonus with rearing and/or falling, barrel roll; 5– repetition of stage 4; 6– generalized tonic-clonic seizure (GTCS), wild running and/or jumping; 7– death). Latency to the first occurrence of each stage and the highest seizure stage reached within 5-minute bins were recorded (*n* = 12-21 per strain, genotype, and sex). We note that one B6.*Scn2a^E/+^* male and one B6.*Scn2a^E/+^* female died shortly after the end of the 120– minute observation period. Statistical comparisons of latency to stage 3, stage 6, and stage 7 were made by log-rank Mantel-Cox time to event analysis. Relevant pairwise comparisons between groups are indicated in **Table 1** (GraphPad Prism).

## Results

### Strain-dependent effects on anxiety-related behavior in Scn2a^E/+^ mice

We used the zero maze and open field assays to assess anxiety-related behavior in adult *Scn2a^E/+^*and WT mice on the B6 and F1D2 backgrounds. Both assays were used as part of our initial characterization of the *Scn2a^K1422E^* mouse model^35^. In the zero maze assay, time spent in the open versus closed arms of the maze was measured (**Figure 1A**). Two-way ANOVA comparing males showed significant main effects of strain and genotype, but no significant strain-by-genotype interaction (**Table 1**). B6.WT and B6.*Scn2a^E/+^* males spent more time in the open arms compared to F1D2 males (**Figure 1A**). The genotype effect was driven by F1D2*.Scn2a^E/+^* males spending significantly more time (12.3 ± 10.7%) in the open arms compared to F1D2.WT males (0.7 ± 1.0%, **Figure 1A**). Two-way ANOVA comparing females also showed significant main effects of strain and genotype, and a significant strain-by-genotype interaction (**Table 1**). B6.WT and B6.*Scn2a^E/+^* females spent more time in the open arms compared to F1D2 females (**Figure 1A**). Also, B6 and F1D2 *Scn2a^E/+^* females spent significantly more time in the open arms compared to B6 and F1D2 WT females. However, the difference between WT and *Scn2a^E/+^* females on the B6 background (23.8%) was larger than the difference between WT and *Scn2a^E/+^*females on the F1D2 background (6.7%, **Figure 1A**). Distance traveled in the zero maze was also measured (**Figure 1B**). Two-way ANOVA comparing males showed significant main effects of strain and genotype, but no significant strain-by-genotype interaction (**Table 1**).

**Figure 1.**
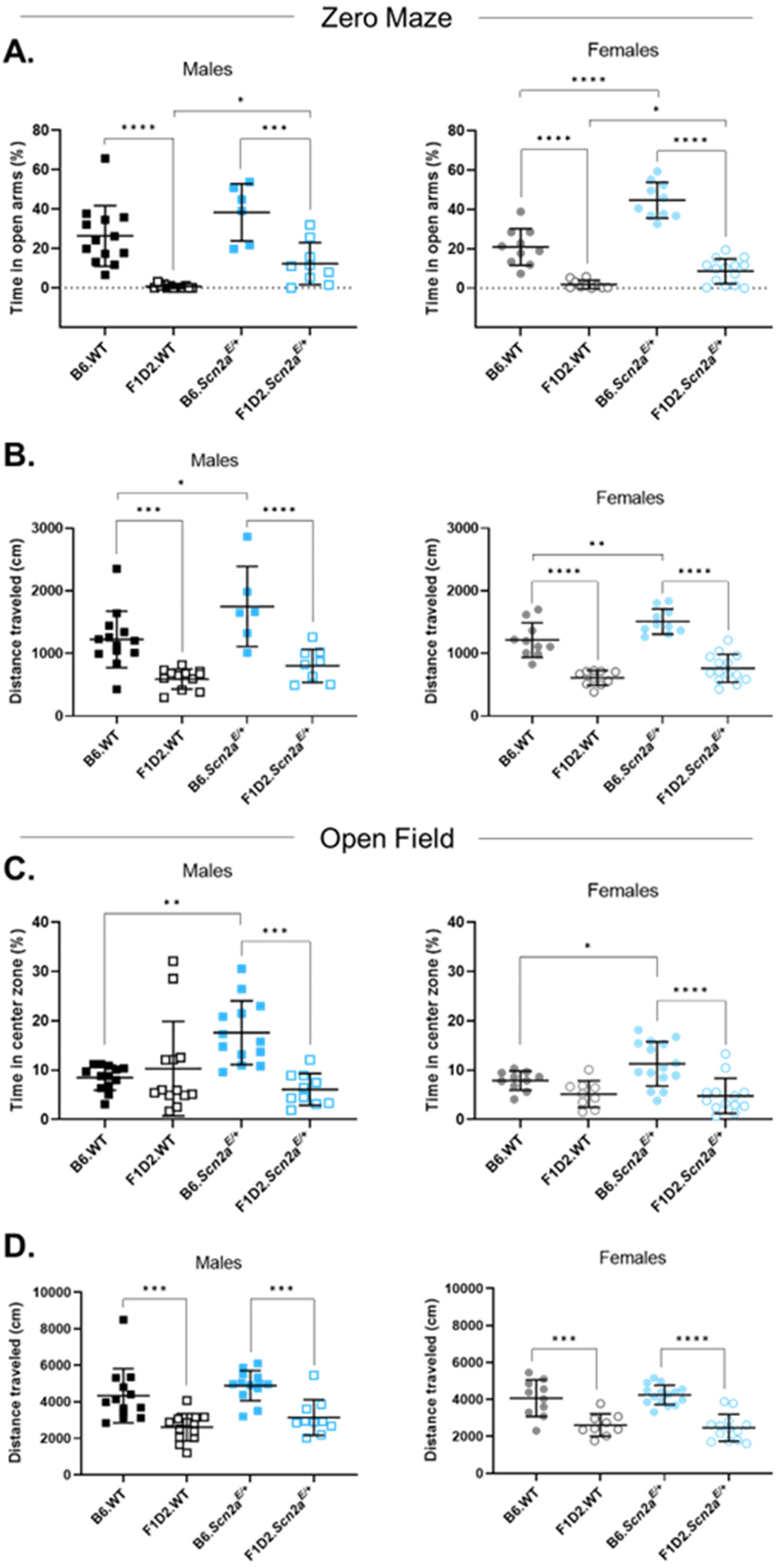
Strain-dependent effects on anxiety-related behavior in *Scn2a^E/+^* mice. (**A**) *Percent time spent in the open arms of a zero maze apparatus.* Two-way ANOVA comparing males showed significant main effects of strain (p<0.0001) and genotype (p=0.0033). B6.*Scn2a^E/+^* and B6.WT males spent more time in the open arms compared to F1D2 males (***p=0.0002; ****p<0.0001). F1D2*.Scn2a^E/+^*males spent more time (12.3 *±* 10.7%), in the open arms compared to F1D2.WT males (0.7 *±* 1.0%; *p=0.0483). Two-way ANOVA comparing females showed significant main effects of strain (p<0.0001) and genotype (p<0.0001) and a significant interaction (p<0.0001). B6.*Scn2a^E/+^* and B6.WT females spent more time in the open arms compared to F1D2 females (*\**\*\*\*p<0.0001; p<0.0001). B6 and F1D2 *Scn2a^E/+^* females spent more time in the open arms compared to B6 and F1D2 WT females (****p<0.0001, *p=0.0163). (**B**) *Total distance traveled in a zero maze apparatus.* Two-way ANOVA comparing males showed significant main effects of strain (p<0.0001) and genotype (p=0.0055). B6.*Scn2a^E/+^* and B6.WT males traveled a greater distance compared to F1D2 males (****p<0.0001; ***p=0.0003). B6.*Scn2a^E/+^* males traveled further (1748 ± 641 cm) compared to B6.WT males (1223 ± 451 cm; *p=0.0163). Two-way ANOVA comparing females showed significant main effects of strain (p<0.0001) and genotype (p=0.0013). B6.*Scn2a^E/+^*and B6.WT females traveled a greater distance compared to F1D2 females (*\*\*\*\**p<0.0001). B6.*Scn2a^E/+^* females traveled further (1508 ± 202 cm) compared to B6.WT females (1213 ± 277 cm; **p=0.0073). (**C**) *Percent time spent in the center zone of an open field arena.* Two-way ANOVA comparing males showed a significant main effect of strain (p=0.0102) and a significant interaction between strain and genotype (p=0.0006). B6.*Scn2a^E/+^* males spent more time (17.6 ± 6.5%) in the center zone compared to F1D2*.Scn2a^E/+^* males (6.1 *± 3.2*%; ***p=0.0002). B6.*Scn2a^E/+^* males spent more time in the center zone compared to B6.WT males (8.5 *±* 2.5%; **p=0.0012). Two-way ANOVA comparing females showed a significant main effect of strain (p<0.0006). B6.*Scn2a^E/+^* females spent more time (11.3 ± 4.5%) in the center zone compared to F1D2*.Scn2a^E/+^* females (4.8 ± 3.6*%;* ****p<0.0001). B6.*Scn2a^E/+^* females spent more time in the center zone compared to B6.WT females (7.9 ± 1.9%; *p=0.044). (**D**) *Total distance traveled in an open field arena.* Two-way ANOVA comparing males showed a significant main effect of strain (p<0.0001). B6.*Scn2a^E/+^* and B6.WT males traveled a greater distance compared to F1D2 males (***p=0.0006; ***p=0.0003). Two-way ANOVA comparing females showed a significant main effect of strain (p<0.0001). B6.*Scn2a^E/+^* and B6.WT females traveled a greater distance compared to F1D2 females (****p<0.0001). For **A-D**, symbols represent individual mice; lines and error bars represent mean ± SD. Males and females were analyzed separately (*n* = 6-13/strain/genotype for males, *n* = 9– 15/strain/genotype for females), using Two-way ANOVA. Displayed p-values calculated by Sidak’s post hoc test.

B6.WT and B6.*Scn2a^E/+^* males traveled a greater distance compared to F1D2 males (**Figure 1B**). The genotype effect was driven by the B6 group. B6*.Scn2a^E/+^* males traveled further (1748 ± 641 cm) compared to B6.WT males (1223 ± 451 cm, **Figure 1B**). Two-way ANOVA comparing females also showed significant main effects of strain and genotype, but no significant strain-by-genotype interaction (**Table 1**). B6.WT and B6.*Scn2a^E/+^*females traveled a greater distance compared to F1D2 (**Figure 1B**). The genotype effect was driven by B6*.Scn2a^E/+^* females traveling further (1508 ± 202 cm) compared to B6.WT females (1213 ± 277 cm, **Figure 1B**).

In the open field assay, time spent in the exposed center zone of the apparatus versus the periphery was measured (**Figure 1C**). Two-way ANOVA comparing males showed a significant main effect of strain and a significant strain-by-genotype interaction (**Table 1**). The strain effect was driven by the *Scn2a^E/+^* group. B6.*Scn2a^E/+^* males spent more time (17.6 ± 6.5%) in the center zone compared to F1D2.*Scn2a^E/+^* males (6.1 ± 3.3%, **Figure 1C**). Furthermore, the strain– by-genotype interaction was driven by the B6 group. B6.*Scn2a^E/+^* males spent more time in the center zone compared to B6.WT males (8.5 ± 2.5%, **Figure 1C**). Two-way ANOVA comparing females showed a significant main effect of strain but no significant effect of genotype or strain– by-genotype interaction (**Table 1**). The strain effect was driven by the *Scn2a^E/+^* group.

B6.*Scn2a^E/+^* females spent more time (11.3 ± 4.5%) in the center zone compared to F1D2.*Scn2a^E/+^*females (4.8 ± 3.6%, **Figure 1C**). Similar to males, B6*.Scn2a^E/+^* females spent more time in the center zone compared to B6.WT females (7.9 ± 1.9%, **Figure 1C**). Distance traveled in the open field was also measured (**Figure 1D**). Two-way ANOVA comparing males showed only a significant main effect of strain, but no significant effect of genotype or strain-by– genotype interaction (**Table 1**). Both B6.WT and B6.*Scn2a^E/+^* males traveled further compared to F1D2 males (**Figure 1D**). Two-way ANOVA comparing females also showed only a significant main effect of strain, but no significant effect of genotype and no significant strain– by-genotype interaction (**Table 1**). B6.WT and B6.*Scn2a^E/+^* females traveled further compared to F1D2 females (**Figure 1D**). The data from these two neurobehavioral assays are generally in agreement with our previous findings that the K1422E variant is associated with lower anxiety–related behavior in mice^35^. In addition, main effects of strain and strain-by-genotype interactions show that the effect of the K1422E variant is more pronounced on the B6 background, thus providing evidence that background strain affects phenotype severity in the *Scn2a^K1422E^* mouse model.

### F1D2.Scn2a^E/+^ mice exhibit rare spontaneous seizures

Previously, we demonstrated that B6.*Scn2a^E/+^* mice exhibit EEG abnormalities, including rare spontaneous seizures without an obvious behavioral component^35^. Therefore, we wanted to investigate whether background strain affects spontaneous seizures in *Scn2a^E/+^* mice. Previous studies have shown that the D2 strain exhibits a high rate of spontaneous spike-wave discharges associated with behavioral arrest and increased susceptibility to induced seizures^36–40^. To evaluate F1D2.*Scn2a^E/+^*mice for spontaneous seizures, juvenile (P28-30) mice were implanted with EEG headmounts for video-EEG monitoring which occurred from 4-7 weeks of age.

F1D2.*Scn2a^E/+^*mice exhibited spontaneous seizures during wake and sleep (**Figure 2**) The seizure shown in **Figure 2** coincided with wild running and abrupt behavioral arrest, whereas seizures in B6.*Scn2a^E/+^* mice occurred only during sleep without a behavioral component. Similar to mice on the B6 background, spontaneous seizures were observed in a minority (2/14) of F1D2.*Scn2a^E/+^*mice. However, similar events were never observed in F1D2.WT mice. The infrequent nature of spontaneous seizures in F1D2.*Scn2a^E/+^* mice precludes a formal analysis of strain-dependence with regard to this particular phenotype. At least qualitatively, it can be said that spontaneous seizures are not more frequent in F1D2.*Scn2a^E/+^* mice compared to B6.*Scn2a^E/+^* mice. However, data from these two strains provide convergent evidence that the K1422E variant is associated with rare spontaneous seizures in mice.

**Figure 2.**
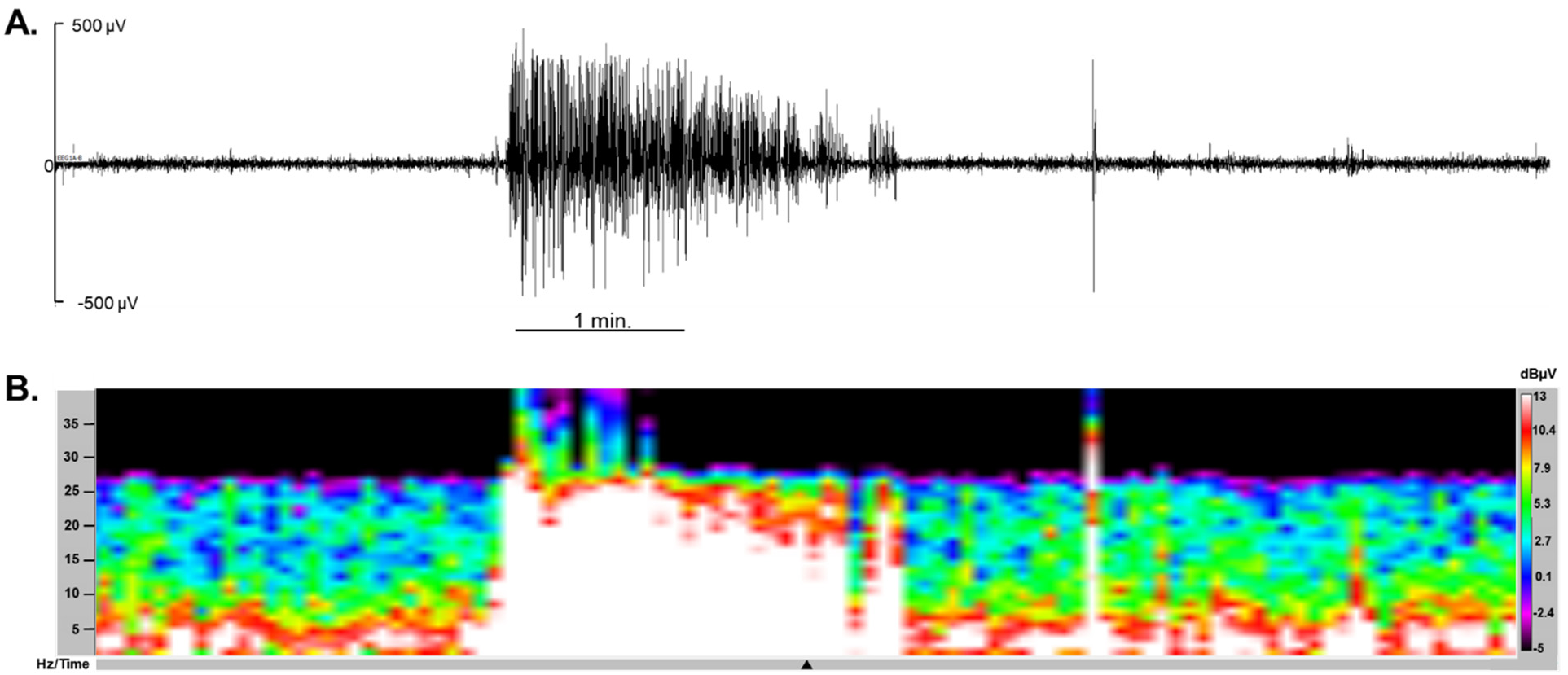
Spontaneous seizures in F1D2.*Scn2a^E/+^* mice. (**A**) Representative 9 min epoch of EEG from F1D2.*Scn2a^E/+^* mice. Signal corresponds to channel 1 (right posterior-left posterior). A localized seizure occurred as an abrupt onset of activity that evolves in amplitude and frequency for ∼2 min before abruptly terminating with return to typical active background. Seizure activity coincided with wild running and abrupt behavioral arrest. (**B**) Spectral density array corresponding to the seizure in (A) showing elevated power (white; dBµV) across the 1-25 Hz frequency range at the time of discharge.

### Susceptibility to KA-induced seizures varies by sex and strain in Scn2a^E/+^ mice

Previously, we demonstrated that B6.*Scn2a^E/+^* mice have a higher threshold for flurothyl-induced seizures compared to WT^35^. We wanted to investigate if similar effects could be observed using a different method of seizure induction and whether background strain affects susceptibility to induced seizures in *Scn2a^E/+^* mice. Previous studies have shown that D2 and commercially available B6D2F1/J mice (analogous to our F1D2 mice obtained by breeding) show greater susceptibility to KA-induced seizures compared to B6 mice^37, 38, 45^. Therefore, we evaluated susceptibility to KA-induced seizures in *Scn2a^E/+^* and WT mice on the B6 and F1D2 backgrounds at 5-6 weeks of age. Seizure severity following KA administration was assessed over a 2 hour period using a modified Racine scale^44^ and the highest stage reached within 5 minute bins was recorded (**Figure 3**).

**Figure 3.**
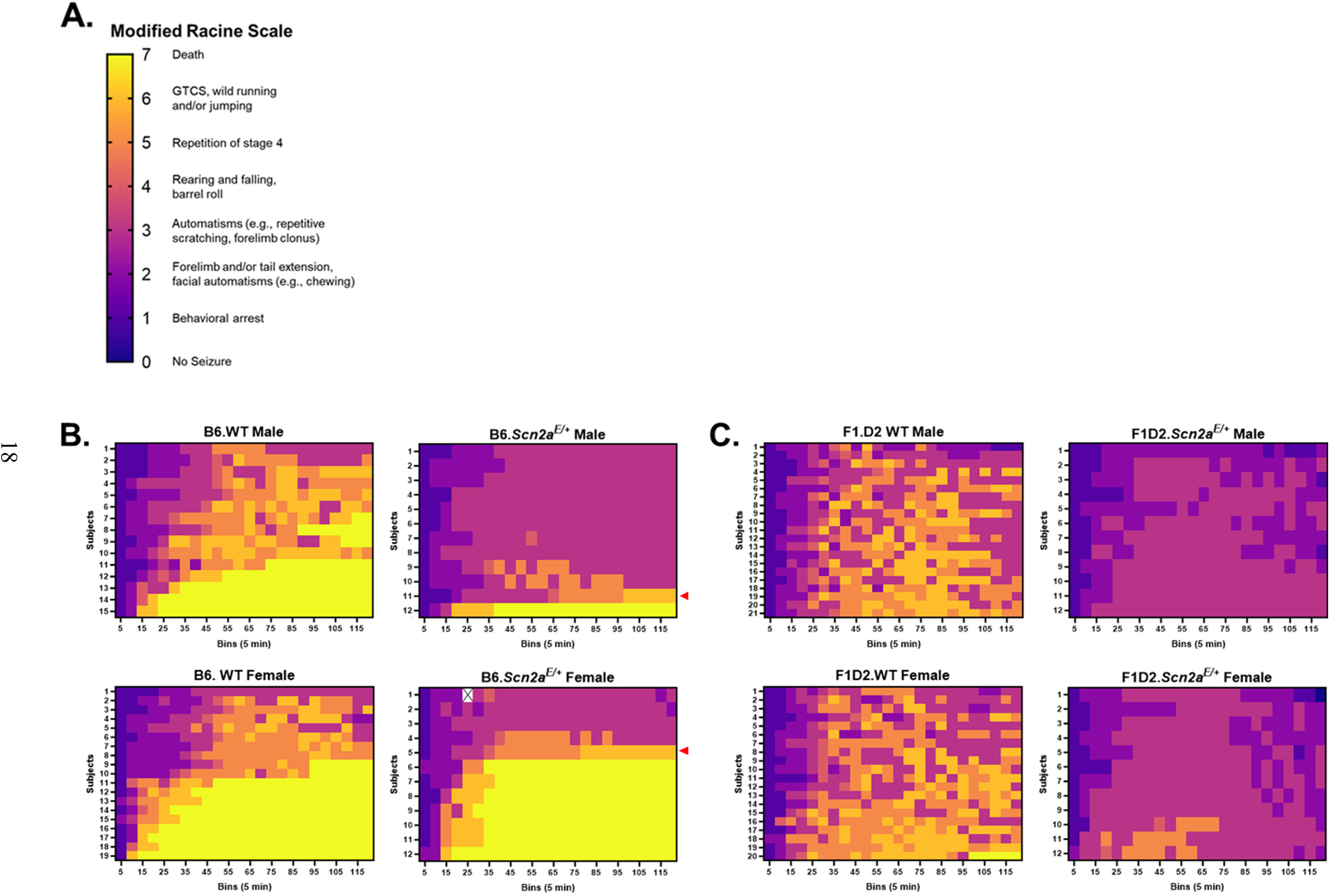
Kainic acid seizure induction in B6 and F1D2.*Scn2a^E/+^* mice. (**A**) Modified Racine scale for scoring seizure severity in mice following KA administration. (**B**) Heat maps depicting individual level seizure severity for mice on the B6 background organized by genotype and sex (*n* = 12-19 per group). Highest seizure stage reached per 5 min time bin was recorded and represented as in (A). Y axis represents individual mice and X axis represents 5 min time bins. A white square with an ‘X’ indicates a 5 min time bin where the subject was obscured during video recording and seizure severity could not be determined. A red triangle denotes a subject that died shortly after the end of the 120-minute observation period. (**C**) Heat maps depicting individual level seizure severity for mice on the F1D2 background organized by genotype and sex (*n* = 12-21 per group). Highest seizure stage reached per 5 min time bin was recorded and represented as in (A). Y axis represents individual mice and X axis represents 5 min time bins.

The heatmaps shown in **Figure 3B** **& C** provide a visual summary of seizure evolution following KA administration for all subjects organized by genotype, strain, and sex. For the purposes of quantitative analysis, we also measured latency to first occurrence of each Racine stage and made pairwise comparisons between groups using by log-rank Mantel-Cox time to event analysis (**Table 1 & Figure 4**). Stages 1-3 of the modified Racine scale represent increasingly severe focal seizure events (**Figure 3A**). Over 99% (122/123) of B6 and F1D2 mice reached stage 3 with minimal differences in latency between groups (**Figure 4A**).F1D2.*Scn2a^E/+^*males reached stage 3 with a median time to event of 25 min compared to F1D2.WT males with a median time to event of 21 min (**Figure 4A**). This suggests that the F1D2.*Scn2a^E/+^* males are less susceptible to seizures induced by KA. We also noted that although 100% (24/24) of B6.*Scn2a^E/+^* males and females reached stage 3, median time to event was longer for males (23 min) compared to females (15 min; **Table 1**). This suggests a sex difference in KA-seizure susceptibility that is both genotype and strain dependent. Stages 4-6 of the modified Racine scale represent increasingly severe generalized seizure events (**Figure 3A**). Significantly fewer B6 and F1D2 *Scn2a*^E/+^ males reached stage 6 compared to strain-matched WT controls. 17% (2/12) of B6.*Scn2a^E/+^*males and 0% (0/12) of F1D2.*Scn2a*^E/+^ males reached stage 6, while 93% (14/15) of B6.WT males and 95% (21/22) of F1D2.WT males reached stage 6 (**Figure 4B**). Interestingly, a genotype difference was only detected in F1D2 females and not B6 females. 0% (0/12) of F1D2.*Scn2a*^E/+^ females reached stage 6 compared to 90% (18/20) of F1D2.WT females (**Figure 4B**). These data provide further evidence to suggest that the K1422E variant is associated with lower KA-seizure susceptibility. We also noted that significantly fewer B6.*Scn2a^E/+^*males, 17% (2/12), reached stage 6 compared to B6.*Scn2a^E/+^* females, 67% (8/12; **Table 1**). Similar to what was observed for latency to stage 3, this suggests a potential sex difference in KA-seizure susceptibility that is both genotype and strain dependent. Stage 7 of the modified Racine scale represents death (**Figure 3A**). Over 98% (64/65) of F1D2 mice did not die following KA administration, with no significant differences based on genotype or sex (**Figure 3C**).

**Figure 4.**
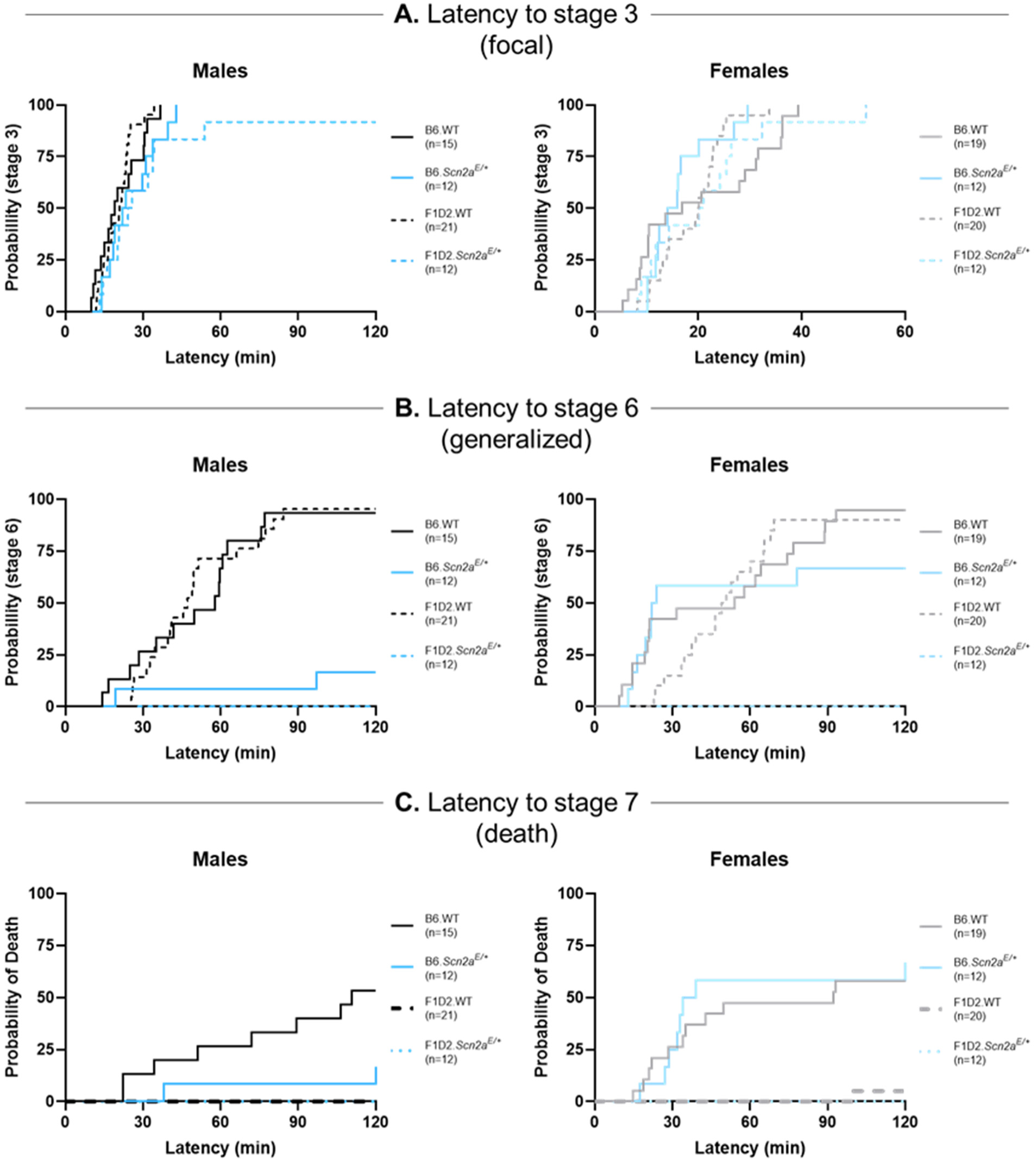
Strain-dependent effects on susceptibility to kainic acid-induced seizures in *Scn2a^E/+^*mice. (**A**) *Latency to first occurrence of modified Racine stage 3 (focal)*. Colored lines represent percentage of subjects that reached stage 3 over time. 99% (122/123) of B6 and F1D2 mice reached stage 3 with minimal in latency between genotype or strain. 92% of F1D2.*Scn2a^E/+^* males reached stage 3 in a median time of 25 min compared to 100% of F1D2.WT males by 21 min (p*=*0.0240). 100% of B6.*Scn2a^E/+^* males and females reached stage 3. The median time to stage 3 was longer (p=0.0073) for B6.*Scn2a^E/+^* males (23 min) compared to B6.*Scn2a^E/+^* females (15 min). (**B**) *Latency to first occurrence of modified Racine stage 6 (GTCS)*. Colored lines represent percentage of subjects that reached stage 6 over time. 17% of B6.*Scn2a^E/+^* and 0% of F1D2.*Scn2a^E/+^*males reached stage 6 compared to 93% of B6.WT and 95% of F1D2.WT males (p<0.0001). Zero F1D2.*Scn2a^E/+^* females reached stage 6 compared to 90% of F1D2.WT females (p<0.0001) and 67% of B6*.Scn2a^E/+^* females (p=0.0008). Fewer B6*.Scn2a^E^*^/+^ males (17%) reached stage 6 compared to B6*.Scn2a^E/+^* females (67%; p=0.0107). (**C**) *Latency to modified Racine stage 7 (death).* Colored lines represent percent mortality over time. 17% of B6.*Scn2a^E/+^* males died post KA administration compared to 53% of B6.WT males (p=0.0467). Zero F1D2.WT males died compared to 53% B6.WT males (p=0.0467). Zero F1D2.*Scn2a^E/+^* females died compared to 67% of B6.*Scn2a^E/+^* females (p=0.0006). 5% of F1D2.WT female mice died compared to 58% of B6.WT females (p=0.0002). Fewer B6.Scn2a^E/+^ males (17%) reached stage 6 compared to B6.Scn2a^E/+^ females (67%; p=0.0077). For **A-C**, males and females are displayed separately but also were compared (*n* = 12-21/strain/genotype for males, *n* = 12–20/strain/genotype for females), using Log-rank Mantel-Cox.

Significantly fewer B6.*Scn2a^E/+^* males, 17% (2/12), died following KA administration compared to B6.WT males, 53% (8/15, **Figure 4C**). Furthermore, zero F1D2.WT males died following KA administration compared to 53% of B6.WT males (**Figure 4C**). As previously noted, a majority of B6.*Scn2a^E/+^* males do not reach stage 6 (i.e. do not have a GTCS), and therefore would not be expected to die as a result of KA-induced status epilepticus. When considering lethality only in those mice that reach stage 6, we observed that both B6.*Scn2a^E/+^* males died following KA administration while approximately half (8/14) of B6.WT males died (Figure **3B**). For females, significantly fewer F1D2 WT and *Scn2a^E/+^* mice died following KA administration compared to genotype-matched B6 controls. 5% (1/20) of F1D2.WT females and 0.0% (0/12) of F1D2.*Scn2a^E/+^* females died, while 58% (11/19) of B6.WT females and 67% (8/12) of B6.*Scn2a^E/+^*females died (**Figure 4C**). However, when considering lethality only in those mice that reach stage 6, we observed that all eight B6.*Scn2a^E/+^* females died following KA administration while 11 of 18 B6.WT females died (Figure **3B**). Furthermore, we noted that significantly fewer B6.*Scn2a^E/+^* males, 17% (2/12) die compared to B6.*Scn2a^E/+^* females, 67% (8/12) when analyzing all lethality (**Table 1**). However, this effect is not observed if we consider group lethality as percentage of subjects that reach stage 6. These data provide evidence to suggest that the K1422E variant is potentially associated with increased risk of death following KA-induced GTCS in mice on the B6 background. Across the three stages analyzed, the results demonstrate that genotype, sex, and strain interact to influence KA seizure susceptibility.

## Discussion

Our rationale for pursuing strain-dependence of phenotypes in the *Scn2a^K1422E^* model was two– fold. First, we wanted to explore the possibility that crossing to a different background strain would unmask more robust phenotypes that would enhance the translatability of the model.

Second, analysis of strain-dependence can serve as the basis for future studies to identify modifier loci and genes. In order to evaluate the effect of genetic background on phenotypes associated with the K1422E variant, we considered alternative mouse strains with endogenous differences in these phenotypes when compared to B6. Relative to B6, D2 mice show higher anxiety-like behavior and higher seizure susceptability^16, 36–40^. Despite these endogenous differences between these two strains, it can be difficult to predict how their respective genetic backgrounds might interact with the pathogenic mechanisms of the K1422E variant to influence disease-related phenotypes. It is similarly difficult to predict how an F1 hybrid background, which is heterozygous for B6 and D2 alleles at all autosomal loci, might interact with the K1422E variant. Thus, our approach to investigating whether background strain affects phenotype severity in the *Scn2a^K1422E^* mouse model is largely exploratory in nature. We showed previously that B6*.Scn2a^E/+^*mice exhibit lower anxiety-like behavior across multiple exploration assays^35^. Using two of the same assays in the current study (zero maze and open field; **Figure 1**), we found that the K1422E variant was associated with lower anxiety-like behavior, consistent with our previous report^35^. In addition, the B6 background was associated with lower anxiety-like behavior relative to F1D2. Furthermore, strain by genotype interactions observed in both assays suggest that lower anxiety-like behavior associated with the K1422E variant is more pronounced on the B6 background (**Figure 1**). These results demonstrate that background strain does affect phenotype severity in the *Scn2a^K1422E^* mouse model.

As part of our initial characterization of B6.*Scn2a^E/+^*mice, we described rare spontaneous seizures without an observable behavioral component^35^. Here, we describe similar seizures in a cohort of F1D2.*Scn2a^E/+^* mice (**Figure 2**). However, some seizures in F1D2.*Scn2a^E/+^* mice were associated with wild running and behavioral arrest, raising the possibility that seizures are more severe on the F1D2 strain compared to B6. Unfortunately, the seizure phenotype shows low penetrance across both strains with only some individuals exhibiting seizures at a rate that precludes quantitative analysis. While certainly more informative than findings from a single strain, the current study only looks at phenotype severity in the context of two genetic backgrounds. A study conducted by Tabbaa & Knoll et al. exemplifies the degree to which genetic background diversity can influence phenotype severity associated with haploinsufficiency in a high confidence ASD risk gene, *Cdh8*^26^. Using panels of genetically diverse recombinant inbred strains from the collaborative cross and C57BL/6J x DBA/2J (BxD) collections, they showed that strain and sex modify penetrance of *Cdh8* haploinsufficiency, with individual strains showing complex profiles of susceptibility and resistance to different traits^26^. Thus, a more thorough examination of phenotypic heterogeneity in the *Scn2a^K1422E^* model using diverse genetic reference panels may reveal a strain with greater susceptibility to spontaneous seizures. At minimum, the observation of spontaneous seizures in F1D2.*Scn2a^E/+^* mice validates our previous findings in B6 mice.

The infrequent nature of spontaneous seizures in B6.*Scn2a^E/+^*mice motivated our investigation of susceptibility to acute, chemically-induced seizures. Previously, we showed that B6.*Scn2a^E/+^*mice exhibit lower susceptibility to flurothyl-induced GTCS^35, 46^. When considering how to approach the question of background strain and seizure susceptibility in the *Scn2a^K1422E^* mouse model, we elected to use an alternative seizure induction paradigm (KA). This would allow us to address whether seizure resistance was inducer-specific, in addition to exploring the effects of background strain. The current study revealed minimal differences between groups in susceptibility to focal seizure events (Racine stage 3; **Figure 4A**) However, *Scn2a^E/+^*mice showed an overall resistance to seizure generalization (Racine stage 6) compared to WT. (**Figure 4B**). This finding is interesting for a few reasons. First, it reminiscent of the resistance to flurothyl seizure generalization we have described in B6.*Scn2a^E/+^* mice, suggesting that this effect is not flurothyl-specific. Next, the resistance to KA seizure generalization associated with the K1422E variant seems to be highly influenced by strain and sex. Nearly 100% of WT mice reached stage 6, regardless of sex or strain compared to 17% of B6.*Scn2a^E/+^* males and 67% of B6.*Scn2a^E/+^* females. Meanwhile, zero F1D2.*Scn2a^E/+^* mice reached stage 6 (**Figure 4B**). This provides strong evidence for the effect of background strain on seizure susceptibility in *Scn2a^E/+^* mice. We also noted that B6.*Scn2a^E/+^* males showed greater resistance to both stage 3 and stage 6 seizures compared to B6 *Scn2a^E/+^* females (the difference in stage 3 was driven by latency rather than occurrence). Importantly, this sex-difference is only observable in *Scn2a^E/+^* mice on the B6 background. One explanation to account for this effect is that modifiers on the B6 background are having a sex-specific effect on phenotype severity associated with the K1422E variant.

Interestingly, a sex-specific effect was observed in B6.*Scn2a^E/+^* mice using the chemoconvulsant flurothyl^35, 46^. Another possibility is that this sex-difference is not specific to B6 and is instead being masked in highly resistant F1D2.*Scn2a^E/+^* mice.

Resistance to seizure generalization associated with the K1422E variant becomes even more compelling when juxtaposed with lethality following KA administration. The data displayed in **Figure 4C** reflect lethality across all subjects included in the study. Interpreted as is, these data suggest that B6.WT males and females as well as B6.*Scn2a^E/+^* females are uniquely susceptible to lethality following KA administration. However, the interpretation changes if lethality is considered as percentage of subjects that die after reaching stage 6. This consideration of lethality is validated by the fact that death in this context is a consequence of a precipitating seizure and no lethality occurred without reaching stage 6 (**Figure 4B**). In this case, 100% of B6.*Scn2a^E/+^* mice died. Given that so few B6.*Scn2a^E/+^* mice reached stage 6, those groups are underpowered relative to others. Similarly, it is not possible to assess the effect of background strain on lethality in this way because no F1D2.*Scn2a^E/+^*mice reached stage 6. Thus, we have preliminary evidence that the K1422E variant is associated with resistance to seizure generalization, but higher risk of death following a convulsive seizure. This further suggests that seizure generalization and lethality are separable phenotypes, which is the subject of a growing body of research^38, 45, 47–49^.

It is estimated that 13% of patients with *SCN2A*-related epilepsy die as a result of sudden unexpected death in epilepsy (SUDEP)^50^. The mechanisms that underlie SUDEP are unclear and likely involve multiple factors but one hypothesis proposes that spreading depolarization in the brainstem mediates cardiorespiratory arrest that leads to death^51–53^. Interestingly, *Scn2a* has been shown to be highly expressed in the mouse brainstem^51, 54^. Moreover, homozygous knockout of *Scn2a* is associated with perinatal lethality due to respiratory depression^51^. We noted similar lethality in mice homozygous for the K1422E variant^35^. These results suggest that *Scn2a* may be important for brainstem mediated control of respiration. The primary biophysical defect associated with the K1422E variant is aberrant calcium influx, which we demonstrated in neurons from *Scn2a^E/+^* mice^35^. This is interesting considering that partial LoF in the gene *Cacna1a* is a protective modifier of lethality in SUDEP-prone *Kcna1* knockout mice^55^. *Cacna1a* encodes the pore forming alpha subunit of P/Q-type calcium channels that mediate neurotransmitter release at terminals and is also associated with epilepsy^56, 57^. Furthermore, GoF mutations in the ryanodine receptor-2 gene that result in “leaky” channels have been linked to SUDEP and have been shown to mediate spreading depolarization in brainstem autonomic microcircuits^58^. Ryanodine receptor-2 is a channel that increases cytoplasmic calcium levels via release from intracellular storage organelles^59^. Based on these findings, it is possible that excess calcium influx through K1422E channels expressed in the nerve terminal could also mediate spreading depolarization in the brainstem following a severe seizure, leading to cardiorespiratory arrest and death. Future mechanistic investigations will explore these possibilities.

Together these data suggest that background strain does affect phenotype severity in the *Scn2a^K1422E^*mouse model. Studies on the effect of genetic background on seizure susceptibility have successfully identified genetic modifiers associated with transgenic models of high risk genes like *Scn2a* and with the endogenously susceptible D2 strain^18, 24, 27–34^. Future studies aimed at identifying modifier loci and genes responsible for the background effects observed here in the *Scn2a^K1422E^* mouse may yield similar results. However, the unique properties of the K1422E variant raise the possibility that unique modifiers may also be identified. Identification of shared modifiers will help expand our understanding of *Scn2a* in health and disease, while the identification of unique modifiers may provide clues regarding the pathological mechanisms of the K1422E variant.

## CRediT authorship contribution statement

**Dennis M. Echevarria-Cooper:** Conceptualization, Methodology, Formal analysis, Investigation, Writing – Original Draft, Writing – Review & Editing, Visualization. **Nicole A. Hawkins:** Conceptualization, Methodology, Investigation, Writing – Review & Editing.

**Jennifer A. Kearney:** Conceptualization, Formal analysis, Writing – Review & Editing, Project administration, Funding acquisition.

## Abbreviations

ASD: autism spectrum disorder
B6: C57BL/6J
D2: DBA/2J
F1D2: [DBA/2JxC57BL/6J] F1 hybrid
GTCS: generalized tonic-clonic seizure
KA: kainic acid
NDD: neurodevelopmental disorders
WT: wild type

## Acknowledgements

We thank Nathan Speakes for technical assistance. This work was supported by the National Institutes of Health grant U54 NS108874 (JAK).

